# Identification of DIR1-dependant cellular responses required for guard cell systemic acquired resistance

**DOI:** 10.1101/2021.06.02.446770

**Authors:** Lisa David, Jianing Kang, Josh Nicklay, Craig Dufrense, Sixue Chen

## Abstract

After localized invasion by bacterial pathogens, systemic acquired resistance (SAR) is induced in uninfected plant tissues, resulting in enhanced defense against a broad range of pathogens. Although SAR requires mobilization of signaling molecules via the plant vasculature, the specific molecular mechanisms remain elusive. The lipid transfer protein-defective in induced resistance 1-1 (DIR1-1) was identified in *Arabidopsis thaliana* by screening for mutants that were defective in SAR. Here we demonstrate that stomatal response to pathogens is altered in systemic leaves by SAR, and this guard cell SAR defense requires DIR1. Using a multi-omics approach, we have determined potential SAR signaling mechanisms specific for guard cells in systemic leaves by profiling metabolite, lipid, and protein differences between guard cells in wild type and *dir1-1* mutant during SAR. We identified two 18C fatty acids and two 16C wax esters as putative SAR-related molecules dependent on DIR1. Proteins and metabolites related to amino acid biosynthesis and response to stimulus were also changed in guard cells of *dir1-1* compared to wild type. Identification of guard cell-specific SAR-related molecules may lead to new avenues of genetic modification/molecular breeding for disease resistant plants.

**One-sentence summary:** DIR1 affects many biological processes in stomatal guard cells during systemic acquired resistance (SAR), as revealed by multi-omics, and it may function through transporting two 18C fatty acids during SAR.

## INTRODUCTION

Since the dawn of agriculture, epidemics of plant pathogens have caused devastating impacts to food production. The plant bacterial pathogen *Pseudomonas syringae* (including more than sixty known host-specific pathovars) infect a broad-ranging and agriculturally relevant plants (Saint-Vincent *et al*., 2020). Although it was first isolated from lilac (*Syringa vulgaris*) in 1899, strains of *P. syringae* are found in many important crops, including beans, peas, tomatoes, and rice (Saint-Vincent *et al*., 2020). *P. syringae* pv tomato (*Pst*) is a pervasive phytopathogenic bacterium that causes damage to a wide range of host crop species. It has been a useful model pathogen for studying host immune response since sequencing and annotation of the 6,397,126 bp genome and two plasmids was funded by the NSF Plant Genome Research Program (Hirano *et al*., 2000). *Pst* infects leaves for chemical nutrients such as carbohydrates, amino acids, organic acids, and ions that are leaked to the leaf apoplast during phloem loading/unloading (Hirano *et al*., 2000). *Pst* causes bacterial brown spot disease in fruit and leaves, damaging crop plants. However, more devastating than brown spot is the unique ability of *Pst* to nucleate supercooled water to form ice. In species of *P. syringe* exhibiting the ice nucleation phenotype, ice-nucleation proteins on the outer membranes of bacterial membranes form aggregates that arrange water into arrays and promote phase change from liquid to solid. The frost-sensitive plants are injured when ice forms in leaf tissues at subzero temperature (Hirano *et al*., 2000). *Pst* has been used extensively to study pathogen infection in numerous host plants including tomato and Arabidopsis. The latter is a reference dicot species with a short life-cycle, fully sequenced genome and rich genetic resources, providing an ideal system to understand how plants may be modified to improve their defense and productivity.

Systemic Acquired Resistance (SAR) is a long-distance plant immune response that improves immunity of systemic tissues after local exposure to a pathogen (Shah *et al*., 2013; David *et al*., 2019). Stomatal pores on leaf surfaces formed by pairs of guard cells are common entry sites for pathogenic bacteria. The specialized guard cells control the opening and closure of stomatal pores in response to environmental conditions (Melotto *et al*., 2008). When stomatal guard cells recognize *Pst* via pattern recognition receptors, stomata close within 1–2 hours and re-open after 3 hours. Re-opening is due to an effector molecule produced by some strains of *P. syringae* called coronatine (COR), which structurally mimics the active form of the plant hormone jasmonic acid-isoleucine (Melotto *et al*., 2008). As a primary entry site for bacteria into the plant tissue, the stomata are at the frontline in plant immune defense (Zhu *et al*., 2012). Our previous research showed that systemic leaves of SAR-induced (“primed”) wild type (WT) Arabidopsis have smaller stomata apertures than control plants, and that *Pst* does not widen stomata aperture in primed leaves, as it does in mock-treated plants (David *et al*., 2020). Reduced stomatal aperture of primed plants correlated with reduced bacterial entry into leaf apoplastic spaces and reduced bacterial proliferation (David *et al*., 2020).

Using a 3-in-1 extraction method to obtain proteins, metabolites and lipids from the same guard cell samples, we conducted multi-omics to identify SAR-related components in guard cells of WT Arabidopsis and a knockout mutant of Defective in Induced Resistance 1 (*DIR1*). *DIR1* encodes a putative apoplastic lipid transfer protein involved in SAR. Arabidopsis plants with mutations in DIR1 exhibit WT-level local resistance to avirulent and virulent *Pst*, but pathogenesis-related gene expression is abolished in uninoculated distant leaves, and mutants fail to develop SAR (Maldonado *et al*., 2002). Champigny *et al*. (2002) examined the presence of DIR1 in petiole exudates from SAR-induced Arabidopsis leaves that were injected with *Pst*. The exudates from the *Pst* injected leaves showed the presence of DIR1 beginning at 30 hour-post-infection (hpi) and peaked at ~ 45 hpi (Champigny *et al*., 2002). Interestingly, the small 7kD DIR1 protein was also detected in dimeric form in the petiole exudates (Champigny *et al*., 2002). DIR1 is conserved in other land plants including tobacco and cucumber, and several identified SAR signals are dependent on DIR1 for long-distance movement, e.g., dehydroabietinal (DA), azelaic acid (AzA) and glycerol-3-phosphate (G3P) (Adam *et al*., 2018). Although most of the LTPs have basic pIs, DIR1 has an acidic pI of 4.25. Martinière *et al*., (2018) found that the apoplastic environment has a more acidic pH than the cellular environment, ranging between 4.0 to 6.3, so perhaps the acidic pI of DIR1 relates to its function in a more acidic environment where it may be neutral, similar to abscisic acid, which is also transported in the apoplast during stress response (Cornish & Zeevaart,1985).

DIR1 is comprised of 77 amino acids, but despite having cystine residues characteristic with lipid transfer proteins (LTP), it has low sequence identity with the previously characterized LTP1 and LTP2 in Arabidopsis. Lascombe *et al*. (2008) compared DIR1 to LTP1 by examining their interactions with various lipid substrates, including lysophosphatidyl cholines (LPCs) with various fatty acid chain lengths (LPC C14, LPC C16, and LPC C18). The results showed that DIR1 showed a greater affinity for LPCs with fatty acid chain lengths with >14 carbon atoms than LTP1. For the LPC with C18 fatty acid tails, the nonpolar C18 end was completely buried within the barrel structure of the DIR1 protein. DIR1 is unique among the LTPs due to its large internal cavity, capable of carrying two lipid molecules, and a proline-rich PxxPxxP motif (including Proline 24 to Proline 30). The Proline-rich regions of DIR1 may be involved in protein-protein interactions, as these regions are located at the surface of the protein and are fully accessible to the aqueous environment (Lascombe *et al*., 2008). These regions are putative candidates for docking of a protein signaling partner, or to other cell components. These features may lend themselves well to its role at a SAR-induced LTP because DIR1 is hypothesized to form a complex with azelaic acid included 1 (AZI1) and localize to the endoplasmic reticulum and plasmodesmata (Yu *et al*., 2013) and to function as a carrier for neutral fatty acids in the apoplast. Many “box-like” LTPs, like DIR1, have a “lid”-like structure that encloses the lipid ligands inside the hydrophobic cavity during transport in the aqueous environment, and have structural motifs that undergo conformational shifts to allow for lipid loading and unloading (Wong *et al*., 2019).

In this study, a multi-omics approach was employed to identify SAR signaling mechanisms specific for stomatal guard cells. The results show potential involvement of DIR1 in amino acid biosynthesis and carbon metabolism in guard cells during SAR. Importantly, four lipid components with long-chain fatty acids were identified as putative DIR1-related SAR signals in guard cells. The results of guard cell molecules in SAR response have not only led to new insights into the basic function of guard cells in the plant immune response, but also may facilitate biotechnology and marker-based breeding for enhanced crop defense.

## RESULTS

### Altered Stomatal Priming Response in *dir1* Correlates with Increased Bacterial Colonization

We have previously characterized that smaller stomata aperture in SAR-induced (primed) WT Arabidopsis plants improves immunity by allowing fewer bacteria to enter apoplastic spaces (David *et al*. 2020). In this study, we examined the role of DIR1 in priming of guard cells during SAR using the *dir1* knockout mutant and its WT ecotype WS. As previously reported for the Arabidopsis Columbia ecotype (Melotto *et al*., 2006; David *et al*. 2021), the basal immune response of the mock-treated WT WS stomata closed after 1 h exposure to *Pst*, and then re-opened after 3 h. In contrast, primed WT WS leaves did not exhibit such stomatal immune responses and maintained a small stomatal aperture during the entire period of *Pst* exposure, similar to that previously observed in the Columbia WT (David *et al*., 2020) (Figure 1A). There was no significant difference in the stomatal aperture from the primed WS leaves taken at 0, 1, and 3 h after *Pst* exposure (Figure 1B). However, guard cells of systemic leaves of *dir1* mutant plants showed an altered response to priming and remain more open at 0 h and 3 h compared to WT. It can be noted that due to the perception of pathogen-associated molecular patterns (PAMPs), the 1 h mock and primed WT and *dir1* apertures are similar. Specifically, average stomatal aperture of primed *dir1* leaves was 1.99 vs. 1.67 μm in WT at 0 h. At 3 h it was 2.80 vs 1.87 μm for *dir1* and WT, respectively. Interestingly, mock-treated *dir1* also showed a larger stomatal aperture at 3 h after exposure to *Pst* when compared to mock-treated WT with an average of 3.60 and 2.69 μm, respectively (Figure 1B).

**Figure 1.**
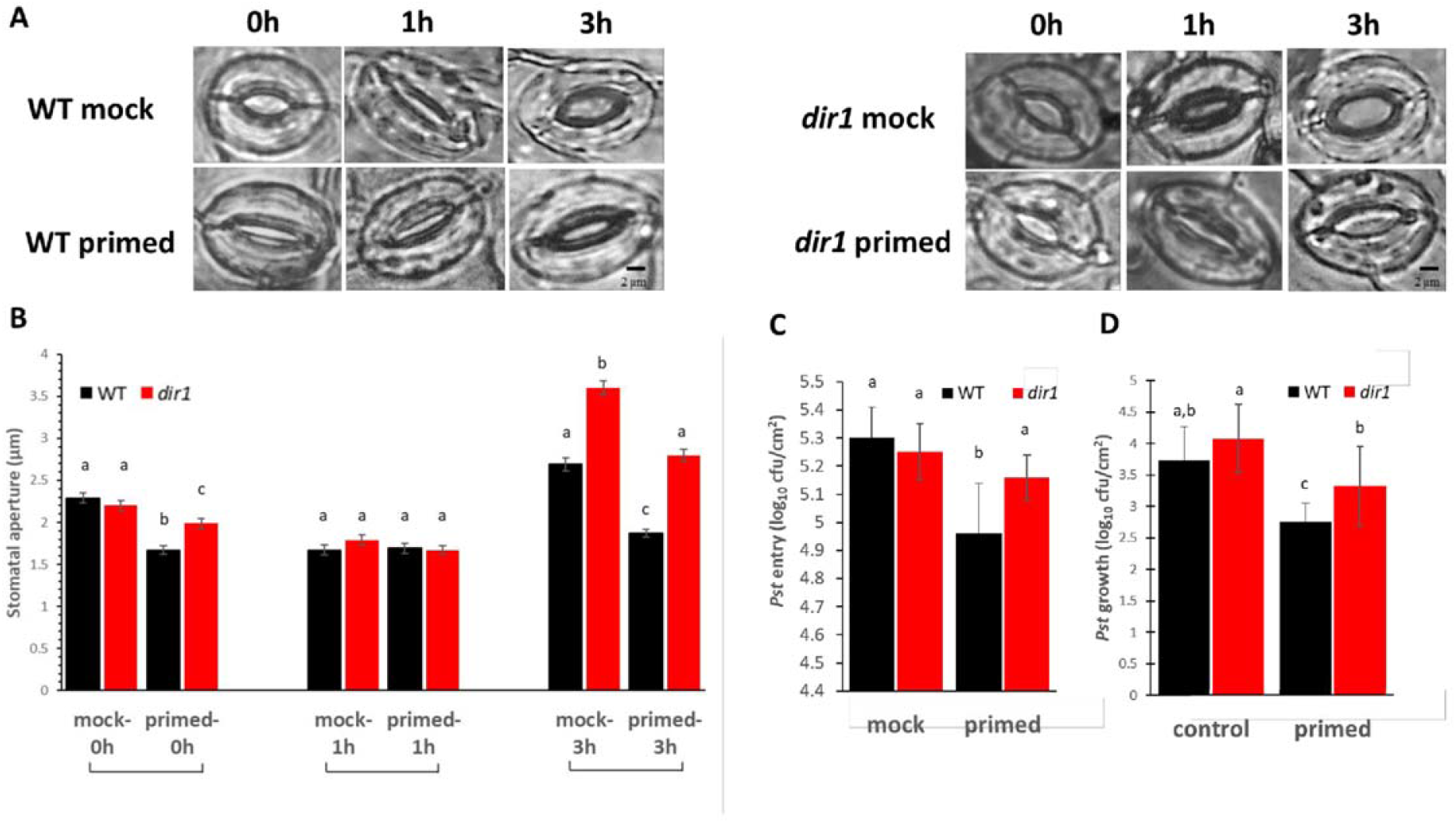
Pathogen entry and growth differences in mock and primed *dir1* mutant and wild type (WT) Arabidopsis leaves. **A.** Images showing representative stomatal apertures in mock and primed *dir1* and WT Arabidopsis leaves after 0, 1, and 3 h after secondary exposure to *Pst* DC3000. **B.** Quantitative measurements of 150 stomata from three replicate experiments. Statistically significant differences were marked by a, b, and c. **C.** *Pst* DC 3000 entry results obtained from nine biological replicates of primed and mock *dir1* and WT plants. The data are presented as average ± standard error. **D.** *Pst* DC 3000 growth results obtained from nine biological replicates of primed and mock plants. The data are presented as average ± standard error with all p-value < 0.05. cfu, colony forming unit.

In the *dir1* mutant, we found that both the control (mock) and primed *dir1* stomatal aperture differed from WT stomata with the same *Pst* treatments (Figure 1). In control plants (mock), we found that the initial (0 h) and PAMP response (1 h) of the *dir1* stomata was not statistically different from that of WT stomata to *Pst* exposure. However, at 3h after exposure to *Pst*, the *dir1* mutant displayed a wider stomatal phenotype, indicating that coronatine secreted from *Pst* had a greater effect on the *dir1* stomata than on the WT (Figure 1A and B). The effect of priming on the stomatal aperture of *dir1* was also different than that of WT. Intriguingly, the *dir1* primed stomata apertures at 0 h were significantly narrower than the control (mock) *dir1* stomata, but less narrow than the WT primed stomata. The mock WT and *dir1* stomata apertures had no significant difference at 0 h (2.29 and 2.20 μm averages, respectively). After priming, WT stomatal aperture decreased to 1.63 μm, but the *dir1* stomata aperture was reduced to only 1.99 μm, making the primed *dir1* stomata apertures significantly different from both the control (mock) *dir1* stomata and the primed WT stomata. The 1 h response to PAMPs from *Pst* was similar regardless of genotype (WT vs. *dir1*) or priming showing the specific response of stomatal closure after PAMP perception. However, at 3 hours post *Pst* treatment, the *dir1* primed stomata phenotype is significantly different from both the *dir1* control and the WT primed. Similar to the stomatal phenotype seen at 0 h, the *dir1* primed stomata had a narrower aperture (2.8 μm) than the *dir1* mock (3.6 μm) but were less narrow than the WT primed (1.87 μm). This demonstrates that although the *dir1* mutant appears to be less resistant to the coronatine from the *Pst* than the WT, it does have improved resistance with priming (Figure 1A and B).

Importantly, the altered stomatal phenotype of *dir1* directly correlates to *Pst* entry into the apoplastic spaces of the leaves and reduced stomatal immunity (Figure 1C). There was no significant difference in the number of *Pst* that were able to enter the apoplast of mock-treated WT, mock-treated *dir1*, or primed *dir1* leaves. Only primed WT stomata were able to reduce *Pst* entry after 3 h exposure to the bacterial pathogen (Figure 1C). Although overall immune response of the *dir1* mutant is reduced, *dir1* plants are still able to mount a SAR response, as demonstrated by the reduced *Pst* growth after 3 days of exposure in the *dir1* primed leaves (Figure 1D).

*Pst* entry and *Pst* growth are not significantly different in the mock-treated *dir1* vs WT plants, correlating to previous evidence that the *dir1* mutant is defective in SAR response, but not in basal pathogen response (Maldonado *et al*., 2002). In primed leaves, stomata apertures correlate to increased *Pst* entry into the apoplast of leaves after 3 hours (Figure 1C) and increased growth of *Pst* after 3 days in the *dir1* mutant when compared to WT (Figure 1D). *Pst* entry is distinctive from *Pst* growth assays because it involves a more rapid time course (within hours after exposure) as opposed to *Pst* growth (measured after 3 days). At 3 h the primed WT plants maintain a smaller aperture upon exposure to *Pst*, while the *dir1* plants have larger stomata apertures. As expected, after 3 h exposure to *Pst*, significantly more bacteria entered the apoplasts in the *dir1* primed leaves compared to the WT primed leaves (Figure 1C). To examine overall susceptibility to *Pst*, we measured bacterial growth in the mock and primed systemic leaves. After 3 days of *Pst* exposure, significantly more bacteria colonized the *dir1* primed leaves than the WT primed leaves. (Figure 1D).

### Differentially Abundant Proteins in the Primed *dir1* and WT Guard Cells

Proteomic analysis of WT versus *dir1* primed guard cell samples taken from distal leaves 3 days after *Pst* treatment identified 2229 proteins, each with more than one unique peptide (1% false discovery rate (FDR)). Of the identified proteins, 155 showed differential abundances in the primed WT guard cells compared to the *dir1* guard cells, with 25 increased in abundance and 130 decreased in abundance, by > 2-fold and a P-value <0.05 (Figure 2A). Of the differentially abundant proteins in *dir1* primed versus (vs) WT primed, only seven were differentially abundant in *dir1*mock vs WT mock, indicating that most changes in protein abundance were due to SAR response, rather than to genotype differences.

**Figure 2.**
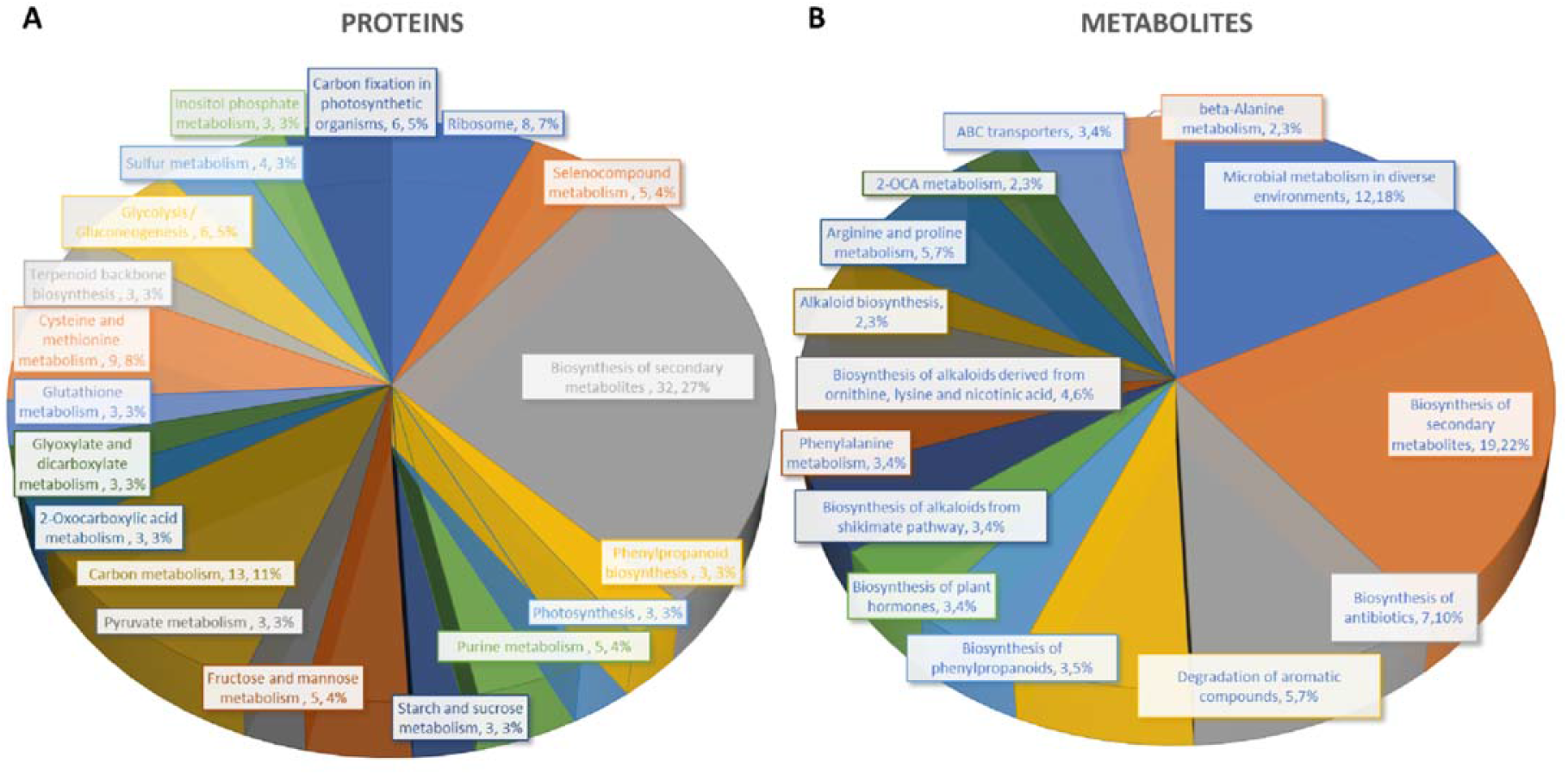
Differential changes of proteins and metabolites in mock and primed *dir1* mutant and WT guard cells. **A.** Biological functions proteins found in KEGG pathways that are differentially abundant in WT versus *dir1* primed guard cells. **B.** Biological functions metabolites found in KEGG pathways that are differentially abundant in WT versus *dir1* primed guard cells.

Of the 155 differential proteins, 76 were mapped to the Arabidopsis KEGG pathway. Again, only three of the 76 were differentially abundant in *dir1* mock vs WT mock. They were phosphoribosylformylglycinamidine cyclo-ligase (mapped to purine metabolism and biosynthesis of secondary metabolites), vacuolar-sorting protein (in endocytosis pathway), and 40S ribosomal protein (in the ribosome pathway). Based on biological functions, the majority of differentially abundant proteins can be broadly categorized into two groups: carbon metabolism-related and amino acid biosynthesis-related. Carbon metabolism-related included 42 proteins from carbon metabolism (13), carbon fixation in photosynthetic organisms (6), glycolysis/gluconeogenesis (6), fructose and mannose metabolism (5), glyoxylate and dicarboxylate metabolism (3), pyruvate metabolism (3), starch and sucrose metabolism (3), and photosynthesis (3). Amino acid biosynthesis-related included cysteine and methionine metabolism (9), and purine metabolism (5). Notably, differentially abundant proteins also grouped into inositol phosphate metabolism (3) related to calcium signaling, terpenoid backbone biosynthesis (3) related to sterols and carotenoids, and glutathione metabolism (3) related to redox signaling (Figure 2A).

Carbon metabolism-related proteins included fructose-bisphosphate aldolase 3 (FBA3), an enzyme involved in the reversible cleavage of fructose-1,6-bisphosphate into dihydroxyacetone phosphate (DHAP) and glyceraldehyde-3-phosphate (GA3P), and two triosephosphate isomerases (TIM and TPI) that catalyze the reversible isomerization between DHAP and GA3P. There three enzymes exhibited 2-fold decreases in the *dir1* primed guard cells compared to WT. Because of the overlap of the carbon metabolism and amino acid biosynthetic KEGG pathways, some differentially abundant proteins were involved in both biological processes, including a pyruvate kinase family protein (PKPα) and an enolase (LOS2). Both were decreased more than 2-fold in *dir1* primed guard cells compared to WT primed (Figure 3).

**Figure 3.**
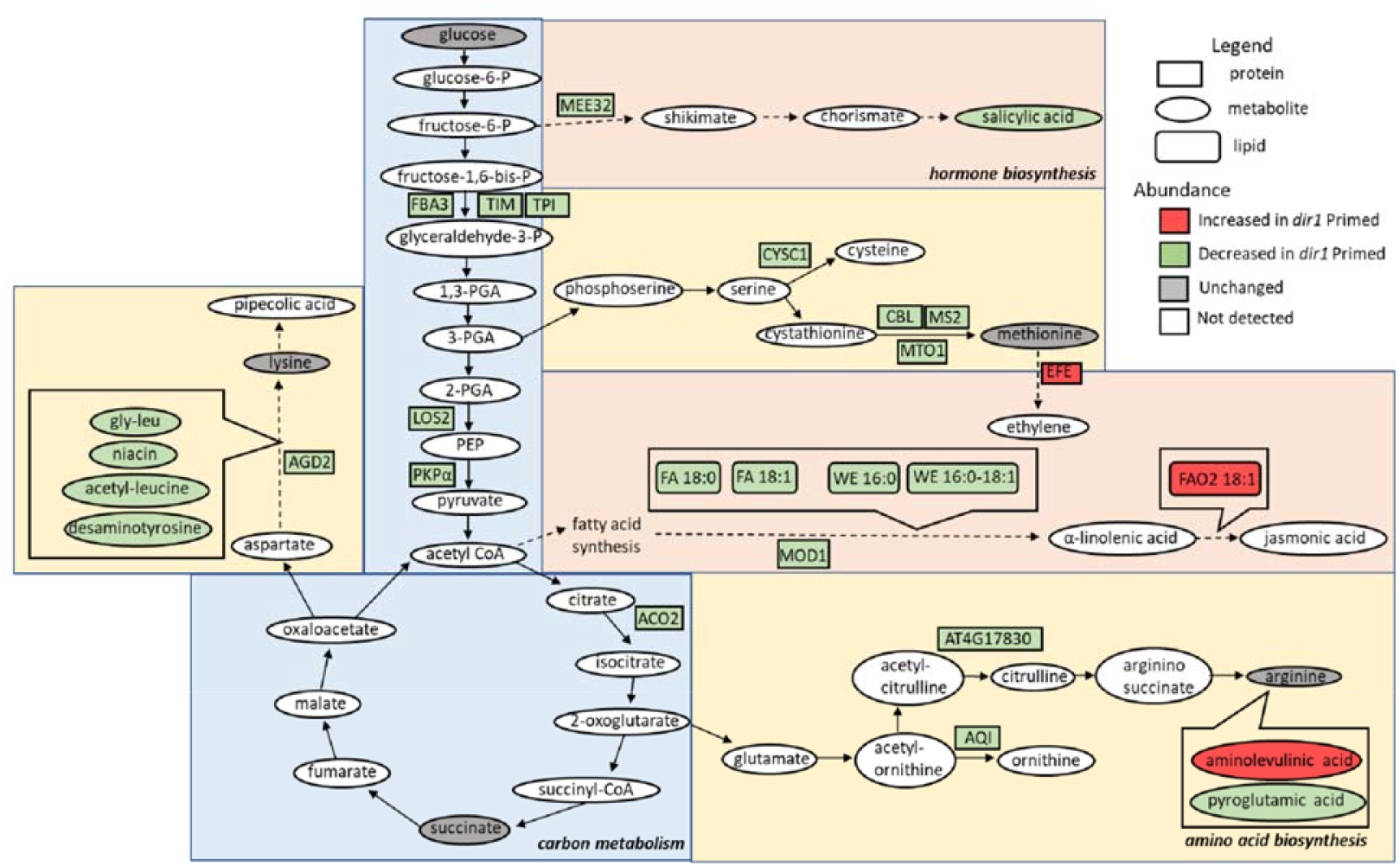
Overview of the role of DIR1 in carbon metabolism, amino acid biosynthesis, and hormone biosynthesis in guard cells during systemic defense response. Loss of *DIR1* results in altered abundance of proteins, metabolites, and lipids involved in carbon metabolism, amino acid biosynthesis, biosynthesis of plant hormones and secondary metabolites. Proteins that were decreased in *dir1* guard cells in the carbon metabolism metabolic pathway included: FBA3, TIM, TPI, LOS2, PKPα, and ACO2. Proteins that were decreased in *dir1* guard cells in the amino acid biosynthesis metabolic pathways included: AGD2, CYSC1, CBL, MS2, MTO1, AT4G17830, and AQI, and decreased metabolites in these pathways included: gly-leu, niacin, acetyl-leucine, desaminotyrosine, and pyroglutamic acid. One increased metabolite in *dir1* guard cells in the arginine biosynthesis pathway was aminolevulinic acid. Proteins that were decreased in *dir1* guard cells in the biosynthesis of hormones and secondary metabolites metabolic pathways included: MEE32 and MOD1, and decreased metabolites and lipids in these pathways included salicylic acid, stearic acid (FA 18:0), behenic acid (FA 18:1), cetyl oleate (WE 16:0/18:1) and ethyl myristate (WE 16:0). One protein, EFE, and one lipid, FAO2 18:1, were increased in these pathways in the *dir1* primed guard cells versus WT primed guard cells. Please refer to Supplemental Table 1 for abbreviations.

The second largest group of differential proteins is related to amino acid metabolism and other pathways with 32 differential proteins between *dir1* vs WT primed guard cells. Some of the proteins are also identified in KEGG biosynthesis of secondary metabolites. For example, Maternal Effect Embryo Arrest 32 (MEE32) is a putative dehydroquinate dehydratase and putative shikimate dehydrogenase. It is found in multiple KEGG pathways including: Biosynthesis of amino acids, Metabolic pathways, Phenylalanine, tyrosine and tryptophan biosynthesis, and Biosynthesis of secondary metabolites. Another example is Aconitase 2 (ACO2) which is also found in multiple KEGG pathways, e.g., Biosynthesis of secondary metabolites, Carbon metabolism, 2-Oxocarboxylic acid metabolism, Glyoxylate and dicarboxylate metabolism, Biosynthesis of amino acids, Citrate cycle (TCA cycle), and Metabolic pathways.

Amino acid biosynthesis-related proteins included aberrant growth and death 2 (AGD2), which encodes a diaminopimelate aminotransferase involved in disease resistance against *Pst* and the lysine biosynthesis via diaminopimelate; methionine synthase 2 (MS2), cysteine synthase C1 (CYSC1) and cystathionine beta-lyase (CBL), which are all involved in cysteine and methionine biosynthesis; and an acetylornithine deacetylase involved in arginine biosynthesis. All mentioned amino acid biosynthesis-related proteins were decreased more than 2-fold in *dir1* primed guard cells compared to WT primed (Figure 3). Differentially abundant proteins involved in redox pathways included glutathione synthetase 2 (GSH2) and glutathione S-transferase TAU 20 (GSTU20) related to redox signaling,

A pathway enrichment analysis was conducted for the differentially abundant proteins using AGRIGO Singular Enrichment Analysis (SEA) (Supplemental Figures S1 and S2). A graphical representation of GO hieratical groups with all statistically significant terms classified levels of enrichment with corresponding colors. The functional enrichment was found in three general groups including response to stimulus, amino acid metabolic processes, and carbohydrate metabolic processes (Supplemental Figure S1). AGRIGO singular enrichment analysis for cellular components revealed enrichment in intracellular organelles including intracellular membrane bounded organelles, plastids, and chloroplast stroma (Supplemental Figure S2).

### Differential metabolites in the primed *dir1* and WT guard cells

A total of 728 metabolites were identified, and 55 metabolites showed significant changes after the priming treatment in the *dir1* versus WT guard cells, with 16 increased and 39 decreased in abundance, by > 2-fold and a P-value <0.05 (Figure 2B). Of these differential metabolites, 34 were mapped to KEGG pathways. When grouping by biological function, the largest group of differentially abundant metabolites found in KEGG pathways were related to biosynthesis of secondary metabolites (19) (Figure 2B).

Several differential metabolites are involved in amino acid biosynthesis and hormone metabolism. For example, SA was decreased by more than 4-fold in the *dir1* primed guard cells compared to WT samples (Figure 3). However, it should be noted that in *dir1* mock versus WT mock the same ratio of decreased SA abundance exits. Metabolites involved in lysine biosynthesis were decreased more than 2-fold in the *dir1* primed guard cells compared to WT. They included gly-leu, niacin, acetyl-leucine, and desaminotyrosine. Metabolites involved in arginine biosynthesis were also changed. For example, pyroglutamic acid that decreased more than 2-fold, and aminolevulinic acid increased more than 4-fold in the *dir1* primed guard cells compared to WT guard cells. Malic acid, which is related to carbon metabolism, was increased 1.8-fold in *dir1* versus WT primed guard cells, but was decreased by nearly 2-fold in *dir1* vs WT mock. When malic acid in the guard cell is pumped out to the apoplast, water moves out reducing turgor pressure in the guard cells and closing the stomata (Santelia and Lawson, 2016).

### Differential Lipids in the Primed *dir1* and WT Guard Cells

A total of 1197 lipids were identified, and 88 lipids showed significant changes in guard cells after the priming of the *dir1* vs WT guard cells (with 37 increased and 49 decreased by > 2-fold). Of the differential lipids, 15 were mapped to KEGG pathways and their biological functions largely fell into two categories: biosynthesis of fatty acids and biosynthesis of secondary metabolites. Notably these lipids included FAO2 18:1, isoleukotoxin diol (DiHOME) involved in linoleic acid metabolism (a precursor for jasmonic acid). It was increased 2.1-fold in the *dir1* vs WT primed guard cells. We also found two long-chain fatty acids (FA) including stearic acid (FA 18:0) and behenic acid (FA 18:1) and two wax esters (WE) including cetyl oleate (WE 16:0/18:1) and ethyl myristate (WE 16:0). They were all decreased more than 2-fold in the *dir1* vs WT guard cells (Figure 3, Figure 4). Ethyl myristate is a long-chain fatty acid ethyl ester resulting from the condensation of the carboxy group of myristic acid with the hydroxy group of ethanol. Palmityl oleate is a wax ester obtained by the condensation of hexadecan-1-ol with oleic acid. Interestingly, both stearic acid and behenic acid were not significantly changed in the *dir1* mock vs WT mock, indicating that this change in FA amount is due to priming, further supporting that they may be the 18C lipid signals potentially transported by DIR1. As to the two wax esters (cetyl oleate and ethyl myristate), they were already more than 2-fold reduced in *dir1* mock vs WT mock, indicating genotypic difference rather than priming effect.

**Figure 4.**
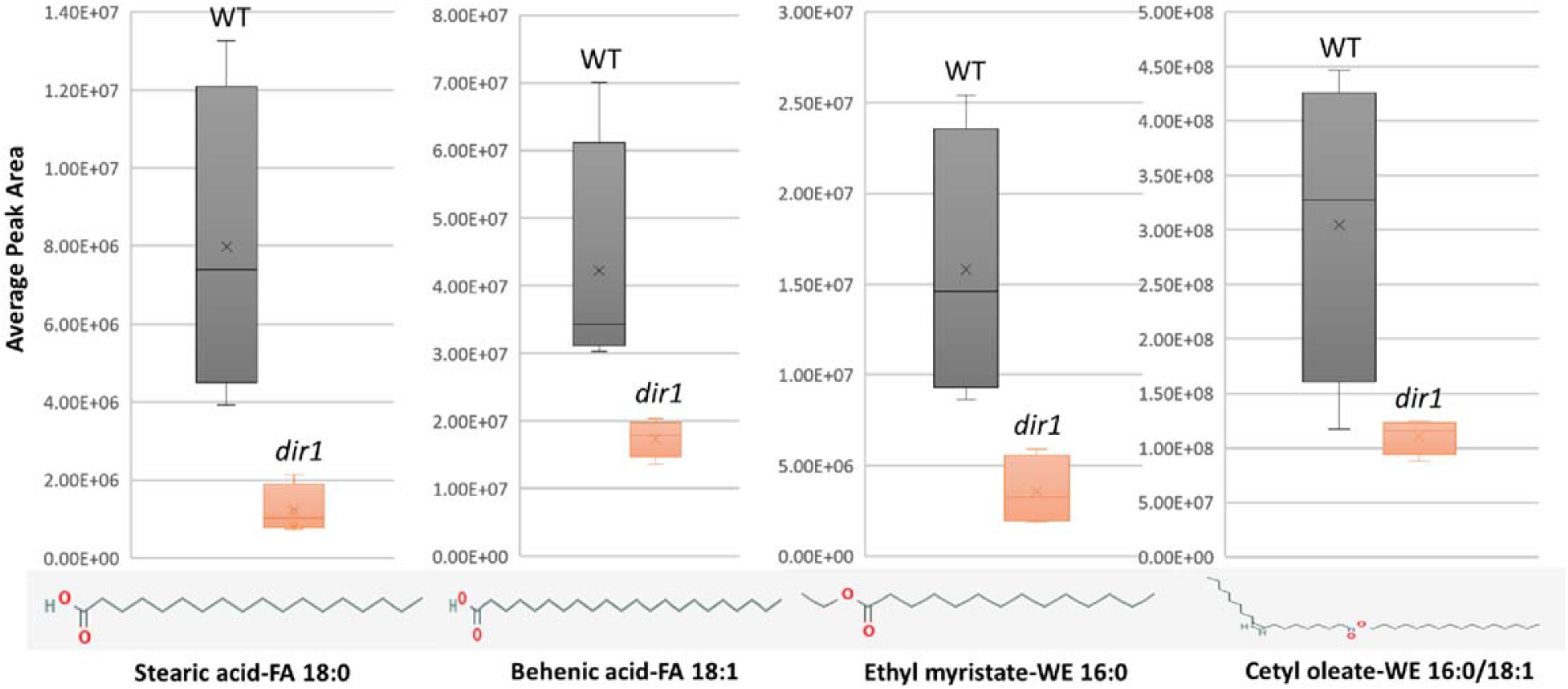
Differentially abundant lipids identified in *dir1* and WT guard cells. **A.** Bar graphs showing decreases of two long-chain fatty acids, stearic acid (FA 18:0) and behenic acid (FA 18:1) and two wax esters, cetyl oleate (WE 16:0/18:1) and ethyl myristate (WE 16:0) decreased > 2-fold in the *dir1* versus WT guard cells. Chemical structures of stearic acid (FA 18:0), behenic acid (FA 18:1), cetyl oleate (WE 16:0/18:1) and ethyl myristate (WE 16:0) are shown. The error bar represents standard deviation of the mean value.

### DIR1 localization and protein interactions with DIR1

Using the Interaction Viewer at the Bio-Analytic Resource for Plant Biology (BAR) (bar.utoronto.ca/eplant), localizations of DIR1 and proteins that interact with DIR1 (AT5G48485) were determined (Figure 5A). Cellular localizations of DIR1 included peroxisomes, Golgi apparatus, endoplasmic reticulum, and plasma membrane. Protein-protein interactions that have been experimentally determined, indicated by the straight, green lines, occur between DIR1 and both ubiquitin-like protein (AT1G68185) and chitin elicitor receptor kinase 1 (CERK1, AT3G21630). Based on Araport 11 annotation, CERK1 is a LysM receptor-like kinase, and has a typical RD signaling domain in its catalytic loop and possesses autophosphorylation activity. GO biological functions of CERK1 include perception and transduction of the chitin oligosaccharide elicitor in innate immune response to fungal pathogens. CERK1 is located in the plasma membrane and cytoplasm and phosphorylates LIK1, an LLR-RLK that is involved in innate immunity (Rebaque *et al*, 2021; Junková *et al*, 2021). However, neither the ubiquitin-like protein nor CERK1 were identified in our proteomics results (Supplemental Table S1).

**Figure 5.**
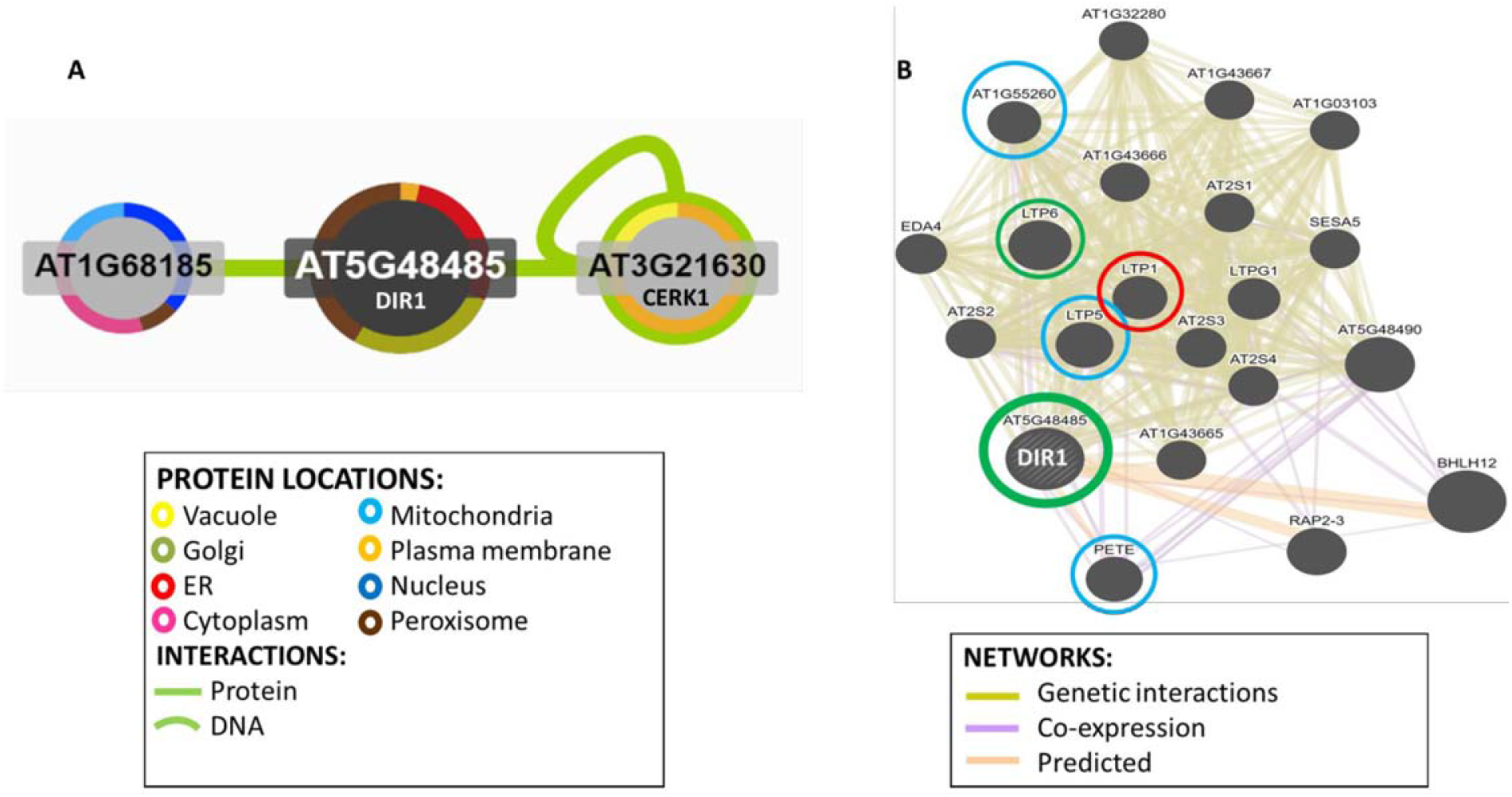
Identification of potential interacting proteins with DIR1. **A.** Protein interaction image was generated using Interaction Viewer at bar.utoronto.ca/eplant. Border color indicates protein location. Green lines indicate protein and DNA interactions that have been experimentally determined. **B.** GeneMANIA tool from bar.utoronto.ca/eplant was used to predict other genes/gene products associated with DIR1 (AT5G48485). Predicted, co-expression, and genetic interaction networks found associated genes/gene products. Proteins identified in guard cell samples are circled. Circle colors indicate increased (red), decreased (green), or unchanged (blue) proteins in *dir1* versus WT primed guard cells.

The GeneMANIA tool at the BAR resource was used to predict other genes/gene products associated with DIR1. Predicted, co-expression, and genetic interaction networks found associated genes/gene products (Figure 5B). In addition to DIR1, our proteomics identified several lipid transfer proteins including LTP1, LTP5, LTP6, Plastocyanin (PETE1) and LTPG6 (AT1G55260) from guard cell samples. LTPG6 is a glycosylphosphatidylinositol-anchored lipid transfer protein involved in defense response to fungus. LTP1 (AT2G38540), is a non-specific lipid transfer protein that binds calmodulin in a Ca^2+^-independent manner. LTP1 is specifically expressed in the L1 epidermal layer and is localized to the cell wall (Fahlberg *et al*., 2019). LTP1, LTP5 (AT3G51600) and LTP6 (AT3G08770) are predicted to encode pathogenesis-related (PR) proteins and are members of the PR-14 protein family (Sels *et al*., 2008). The mRNA of LTP1 is cell-to-cell mobile (Bogdanov *et al*., 2016). PETE1 is one of two Arabidopsis plastocyanins (PETE1 and PETE2). Its mRNA expression is one-tenth of the level of *PETE2*. Although PETE2 is involved in copper homeostasis, PETE1 is not responsive to increased copper levels, but it may participate in electron transport during copper-limiting conditions (Abdel-Ghany, 2009; Weigel *et al*., 2003). DIR1 was not present in our *dir1* knockout mutant samples, and LTP6 was significantly decreased in the *dir1* versus WT after priming. LTP1 was increased in *dir1* versus WT, and LTP5, PETE and LTPG6 were unchanged during priming in *dir1* versus WT guard cells (Figure 5B). DIR1 was associated with LTP1, LTP5, LTP6 and LTPG6 via genetic interaction networks, and with PETE1 via predicted and co-expression networks (Figure 5B).

## DISCUSSION

### *dir1* is Deficient in Both Local and Systemic Guard Cell Immune Responses

Although SAR has largely been studied at the level of leaf or whole plant level, we have recently shown evidence that SAR affects guard cell response to the bacterial pathogen *Pst* (David *et al*., 2020). DIR1 is required for movement of several chemically diverse SAR signals including DA, G3P, AzA, and possibly MeSA (Adam *et al*., 2018). As we have recently reported stomatal movement and guard cell molecular changes underlying stomatal SAR responses (David *et al*., 2020), here we first characterized the stomatal movement phenotype of the *dir1* mutant versus WT in response to *Pst*. Results from our work and previous studies (Melotto *et al*., 2008; Pang *et al*., 2020) clearly showed that stomatal guard cells from different genotypes of Arabidopsis (WS and Columbia) exhibited similar basal immune responses. After priming for three days, stomata from the WT WS leaves had an initial narrow aperture compared to control (mock) stomata, and they maintain this narrow aperture during PAMP perceptions at 1 hour and also at 3 hours after *Pst* treatment. This result is also similar to the Columbia WT plants (David *et al*., 2020),

In the *dir1* mutant, at 3h after exposure to *Pst* the *dir1* mutant displayed a larger stomatal aperture, indicating that coronatine secreted from *Pst* had a greater effect on the *dir1* guard cells than on the WT. The effect of priming on the *dir1* stomata was also different from the WT stomata. The *dir1* primed stomata apertures at 0 h were narrower than the mock *dir1*, but less narrow than the WT primed stomata. At 3 h post *Pst* treatment, the *dir1* primed stomatal aperture is smaller than mock-treated, but less narrow than the WT primed. The altered stomatal aperture of *dir1* directly correlates to *Pst* entry into the apoplastic space (Figure 1). Clearly, although the *dir1* mutant appears to be less resistant to the coronatine than the WT, it does have improved resistance after priming. This result is consistent with previous literature, which showed a partial SAR-competent phenotype of *dir1* (Champigny *et al*., 2013). Although the partial SAR-competent phenotype of *dir1* was able to reduce the *Pst* growth, it did not decrease the entry of *Pst* via the stomatal pores. Therefore, *dir1* is deficient in both local and systemic guard cell immunity.

### DIR1 affects Guard Cell Carbon Metabolism and Amino Acid Biosynthesis During SAR

Based on our multi-omic results, the majority of the differential proteins and metabolites (including lipids) were in the carbon metabolism, amino acid biosynthesis and secondary metabolite biosynthesis pathways (Figures 2 and 4). Most of the molecules were lower in the primed *dir1* guard cells than the primed WT guard cells. These results indicate that DIR1-dependent SAR is necessary for regulation of amino acid biosynthesis and secondary metabolites in guard cells. It also indicates that guard cells attenuate their carbon metabolic pathways to divert resources to amino acid biosynthesis in response to priming in WT, and that this process is at least partially dependent on DIR1 in guard cells. In addition, the differential proteins enriched for plastid and chloroplast components again support alterations in carbon metabolic pathways induced by SAR.

One interesting aspect of our results is that we did not identify changes in pathogenesis-related (PR) proteins in the *dir1* primed guard cells. Similarly, the abundance of AzA was not significantly different in the primed *dir1* versus WT guard cells. On the other hand, the key regulatory SAR metabolite SA showed a 50-fold decrease in the primed *dir1* vs WT guard cells. Previously, we reported that primed guard cells in uninoculated leaves of Arabidopsis narrowed stomatal apertures, reduced entry of *Pst* into the leaves, and had increased SA in primed guard cells compared to mock guard cells (David *et al*., 2020). The lower SA in the primed *dir1* guard cells correlates well with our previous findings and demonstrates that DIR1 is required to transmit the long-distance SAR signal to the guard cells in uninfected leaves and increase SA in the primed guard cells. Recently, translocation of SA from primary infected tissue to distal uninfected leaves was shown to likely occur via the apoplastic space between the cell wall and plasma membrane (Lim *et al*., 2016). Unlike the SAR-induced signals G3P and AzA, which were preferentially transported via symplastic transport and through plasmodesmata, pathogen infection resulted in increased SA accumulation in the apoplastic compartment, and SAR-induced accumulation was unaffected by defects in symplastic transport via plasmodesmata (Lim *et al*., 2016). Mature guard cells have callose depositions that block plasmodesmata, and thus SAR-signals that can be transported via the apoplast, rather than the symplast, would logically be able to affect the guard cells during SAR. Alternatively, SA could be *de novo* synthesized in the primed guard cells. This SA biosynthesis is also affected by *DIR1* mutation. How DIR1 regulates SA biosynthesis is not known.

### Lipidomics Revealed Four Long-Chain Fatty Acids Associated with DIR1

Previously, we found that fatty acids were increased in the primed WT guard cells (David *et al*., 2020). Here we compared the levels of lipids found in primed WT guard cells to those in the *dir1* mutant. Our goal was to identify lipids in guard cells that are dependent on DIR1 during priming. DIR1 has been characterized as a lipid transfer protein, and the core of its structure forms a left-handed super helical arrangement of four α-helices building the hydrophobic central cavity. Lascombe *et al*. (2008) demonstrated that DIR1 showed a greater affinity for LPCs with fatty acid chain lengths with >14 carbon atoms and that nonpolar C18 fatty acid tails were completely buried within the barrel structure of the DIR1 protein, presumably allowing non-polar fatty acids to be transported in polar cellular environments. The two long-chain fatty acids (stearic acid (FA 18:0) and behenic acid (FA 18:1)) and two wax esters (cetyl oleate (WE 16:0/18:1) and ethyl myristate (WE 16:0)) were all decreased > 2-fold in the *dir1* guard cells compared to WT guard cells (Figures 3 and 4). As both stearic acid and behenic acid were not significantly changed in *dir1* mock vs WT mock, this change in FA levels is likely due to priming, further supporting that they may be the 18C lipid signals transported by the DIR1. Further analysis is required to determine the relationship between DIR1 and these 18C fatty acids. It is reasonable to propose that DIR1 may transfer stearic and behenic acid to guard cells during SAR. Previously we identified an increase in palmitic acid and its derivative 9-(palmitoyloxy) octadecanoic acid in primed WT guard cells and proposed that fatty acids could allow for the development of lipid rafts or other alterations of membrane structure in guard cells, modulating stomatal immune responses (David *et al*., 2020).

Plant wax esters are neutral lipids with long-chain (C_16_ and C_18_) or very-long-chain (C_20_ and longer) carbon structures and are mostly found in cuticles where they provide a hydrophobic coating to shoot surfaces (Li *et al*., 2008). Recently the cuticle has been shown to regulate transport of SA from pathogen-infected to uninfected parts of the plant via the apoplast during SAR (Lim *et al*., 2020). Lim *et al*. (2020) found that cuticle-defective mutants with increased transpiration and larger stomatal apertures reduced apoplastic transport of SA and caused defective SAR response. It is interesting to note that our results demonstrate that WT stomata maintain narrow stomata apertures after priming, potentially to reduce transpiration and increase water potential, and possibly routing SA to the apoplast. The *dir1* mutant, on the other hand, had larger stomatal apertures, perhaps resulting in defect in SA movement in the apoplast. It is not known whether the mutant has defect in cuticle structure due to the decreases of wax esters (cetyl oleate and ethyl myristate). However, since the decreased cetyl oleate and ethyl myristate in *dir1* guard cells after priming were already >2-fold reduced in *dir1* mock vs WT mock, this was a genotypic difference, rather than a result of priming. If, as reported by Lim *et al*. (2020), defects in the cuticle reduce transport of SA, the reduced wax esters in *dir1* vs WT could explain the reduce SA in *dir1* guard cells (both mock and primed) and contribute to the SAR defect of the *dir1* mutant.

One cuticle-defective mutant was a knockout of *MOD1*, an acyl carrier protein (ACP) which transports a growing FA chain between enzyme domains of FA synthase during FA biosynthesis. The *mod1* mutant is defective in the key FA biosynthetic enzyme enoylACP reductase and has reduced levels of multiple FA species and total lipids (Lim *et al*., 2020). Interestingly, we also found that MOD1 was > 2-fold lower in *dir1* guard cells versus WT guard cells after priming (Figure 3). This result supports our previous results that FA synthesis plays a key role in SAR priming in guard cells (David *et al*., 2020). However, how DIR1 affects MOD1 and FA biosynthesis awaits further investigation.

## CONCLUSION

Guard cells that control stomatal aperture respond to various abiotic and biotic signals and have membrane-bound pattern recognition receptors that perceive bacterial pathogens. One neglected area of SAR research has been the role that stomatal guard cells play in SAR. This work investigates the role of SAR-related lipid transfer protein DIR1 in guard cell-specific SAR. After priming and also after exposure to the bacterial pathogen *Pst*, stomata of WT remain at a narrow aperature. In contrast, the *dir1* mutant showed defects in stomatal closure. Based on the multi-omics data, proteins and metabolites related to amino acid biosynthesis, secondary metabolism and response to stimulus were altered in guard cells of *dir1* compared to WT. For example, several proteins in the methionine biosynthesis pathway and a protein related to ethylene biosynthesis were decreased in the *dir1* primed guard cells compared to WT. It is known that ethylene is biosynthesized via methionine and ethylene plays a role in SA-regulated stomatal closure by mediating ROS and nitric oxide (Wang *et al*., 2020). A putative shikimate dehydrogenase was also decreased in the *dir1* guard cells after priming. As SA is a product of the shikimate pathway and was also lower in *dir1* guard cells, the slow-down in this pathway could explain the decrease of stomatal closure and defense observed in the *dir1* mutant during priming. Our lipidomics results highlighted a role for fatty acid signaling and cuticle wax esters in the primed guard cells, i.e., two 18C fatty acids as putative lipid mobile signals and two 16C wax esters dependent on DIR1. These results are also correlated to a decrease in the MOD1 in the *dir1* guard cells. As *mod1* mutants have been shown to have cuticle defects and reduced transport of SA to distal tissue during SAR, this relates to the decreased SA in the *dir1* guard cells. Multi-omics has shown utility in discovering DIR1-dependent molecular networks in stomatal immunity. The improved knowledge may facilitate effort in biotechnology and marker-based breeding for enhanced plant disease resistance.

## MATERIALS AND METHODS

### Plant Growth and Bacterial Culture

*A. thaliana* WS seeds were obtained from Arabidopsis Biological Research Center (Ohio, USA). They were suspended in deionized H_2_O and vernalized at 4 °C for two days before planting. The seeds were cultivated in soil and grown in controlled environmental chambers in short day (8-hour light/16-hour dark) environment. The temperatures during the light and dark periods were 22 °C and 20°C, respectively. Incandescent bulbs capable of emitting 140 μmol m^-2^ s^-1^ at the leaf surface were used in the growth chamber with a relative humidity of about 60%. A dome was placed over the flat until seeds began germination. After 2 weeks of growth, seedlings were transferred into individual pots. Plants were watered weekly, kept in the chamber until mature rosette (stage 3.9), and observed at 5-weeks of age.

*Pseudomonas syringae* pv. *tomato* DC3000, the model pathogen for Arabidopsis SAR induction was used for the experiments. Agar media plates were made using King’s B media protocol. A 1-liter solution contained 20 g Protease peptone No. 3, 1.5 g K_2_HPO_4_ (s), 0.75 g MgSO_4_ (s), 10 mL glycerol, 15 g agar, and deionized H_2_O King’s B Media was autoclaved, and antibiotics Rifampicin (25 mg/L) and Kanamycin (50 mg/L) were added once the solution is cooled. Solution with agar was used for plates and *Pst* colonies were streaked on this medium and incubated for overnight at 28 °C. *Pst* colonies were grown in the same King’s B media without agar in solution overnight, pelleted by centrifugation at 6000x g for 10 min, and used for treatment of Arabidopsis plants.

### Stomata Aperture Measurements

Primary inoculation occurred via needless syringe infiltration, where the leaves were either primed with *Pst* DC3000 (OD_600_= 0.02) suspended in 10 mM MgCl_2_ or mock-treated with 10 mM MgCl_2_. At 3 days post inoculation, the leaf opposite to the injected leaf was detached for a secondary treatment. In the secondary treatment, the leaves were either floated in 10 mM MgCl_2_ or in *Pst* DC3000 (OD_600_= 0.2, in 10 mM MgCl_2_) in small petri-dishes. Three leaves were used for each time point and secondary treatment group, and only one leaf was collected from each plant. Stomatal apertures were measured at three time points: 0 h, 1 h and 3 h. The leaves were collected and peeled using clear tape. The peel from abaxial side of the leaf was then placed on a microscope slide and images were collected using a DM6000B light microscope (Leica, Buffalo Grove, IL USA) This experiment was repeated 3 times to image 50 stomata from each replicated treatment and a total of 150 stomata measurements from 3 independent replicates were analyzed for each time point. Stomatal apertures were measured using ImageJ software (National Institutes of Health, Bethesda, MD, USA, (http://imagej.nih.gov/ij/).

### *Pst* DC3000 Entry and Growth Assays

To measure how much bacteria entered the apoplast after three hours, nine plants from three independent experiments were grown to 5-weeks and prime-treated via infiltration with either *Pst* DC3000 (OD_600_= 0.02) or mock-treated with 10 mM MgCl_2_. Three days later, the leaf opposite to the one infected was detached and floated in *Pst* (OD_600_= 0.2) solution for both mock and primed plants. After 3 h, the leaf was placed in a Falcon tube with 0.02% Silwet (Su *et al*, 2017), vortexed for 10 seconds, dried with sterile Kim wipes, wrapped in clean aluminum foil, and taken to Laminar flow hood for aseptic treatment. In the hood, an autoclaved hole-puncher was used to obtain one disk from each leaf (0.5 cm diameter), and the disk was placed in 100 μL sterile H_2_O. Each leaf disk was then ground using an autoclaved plastic grinding tip, and 10 μL of the solution was collected to make a 1:1000 serial dilution. From the dilution, 100 μL was collected and plated on agar media containing Rifampicin (25 mg/L) and Kanamycin (50 mg/L). After 2 days of incubation at 28 °C, the colonies on the plate were counted. The experiment was done 3 times with 3 replicates each time. The bacterial counts of nine replicates were used to calculate mean and standard error.

*Pst* growth experiment determines how much bacteria grow in the apoplast after 3 days. *Arabidopsis* plants (9 independent replicates) were grown to 5-weeks and either mock or prime-treated. After three days of treatment, the rosette leaves were sprayed with *Pst* DC3000 (OD_600_= 0.2) and a dome was put on top for 24 hours. After 24 hours, the dome was removed, and the plants were left in growth chamber for another 48 hours. One opposite leaf of each plant was then detached, washed in 0.02% Silwet, and one disk was taken from leaf to make a 1:1000 serial dilution and plate it on media. Colonies were counted to determine how much bacteria were able to grow in the apoplast. The experiment was repeated 3 times with 3 replicates each time. The bacterial counts from the nine replicates were used to calculate mean and standard error.

### Isolation of Enriched Guard Cells for Multi-omics Experiments

Enriched guard cell samples were prepared as described in Kang *et al* (2021). Briefly, for each sample 144 mature leaves were collected from 36 individual plants. After removing the midvein with a scalpel, the leaves were blended for 1 minute in a high-speed blender with 250 mL of deionized water and ice. The sample was then filtered through a 200 μm mesh filter. This process was repeated 3 times to obtain intact stomatal guard cells, which were collected immediately into 15 mL Falcon tubes, snap frozen in liquid nitrogen, and stored in −80 °C. Guard cell viability and purity was verified by staining with fluorescein diacetate and neutral red dye, which showed that guard cells remained intact and viable.

### 3-in-1 Extraction of Proteins, Metabolites and Lipids from Guard Cell Samples

We adapted a protocol to simultaneously extract metabolites, lipids, and proteins from a single whole leaf or guard cell sample (Kang *et al*., 2021). Briefly, a chloroform and methanol solution is added to samples that are in an aqueous isopropanol solution. This process induces the formation of two solvent layers – an upper aqueous phase containing hydrophilic metabolites, and a lower organic phase containing lipids and other hydrophobic metabolites. The proteins are at the interphase. Components were normalized from internal standards that were added during the first step of extraction. Internal standards included, for proteins: 60 fmol digested bovine serum albumin (BSA) peptides per 1 μg sample protein; for metabolites: 10 μL 0.1 nmol/μL lidocaine and camphorsulfonic acid; and for lipids: 10 μL 0.2 μg/μL deuterium labeled 15:0–18:1(d7) phosphatidylethanolamine (PE) and 15:0–18:1(d7) diacylglycerol (DG). The lipid extracts were dried under nitrogen gas to prevent oxidation and stored in −80 °C. The lipid extract was later dissolved in 1 mL isopropanol for LC-MS/MS analysis. Aqueous metabolites were lyophilized and placed at −80 °C. Aqueous metabolite pellets were later solubilized in 100 μL 0.1% formic acid for LC-MS/MS analysis. Protein components were collected by precipitation in cold 80% acetone in the centrifuge tubes at −20 °C overnight. Acetone was removed using glass pipettes, and the tubes with the protein samples were dried in a speedvac.

### Protein Digestion and LC-MS/MS

Four biological replicates of mock and SAR primed guard cell samples from WT and *dir1* genotypes were prepared for proteomic experiments. Protein samples were resuspended in 50 mM ammonium bicarbonate, reduced using 10 mM dithiothreitol (DTT) at 22 °C for 1 h, and alkylated with 55 mM chloroacetamide in darkness for 1 h. Trypsin (Promega, Fitchburg, WI) was added for digestion (w/w for enzyme: sample = 1: 100) at 37 °C for 16 h. The digested peptides were desalted using micro ZipTip mini-reverse phase (Millipore), and then lyophilized to dryness. The peptides were resuspended in 0.1% formic acid for mass spectrometric analysis.

The bottom-up proteomics data acquisition was performed on an EASY-nLC 1200 ultraperformance liquid chromatography system (Thermo Scientific) connected to an Orbitrap Exploris 480 with FAIMS Pro instrument (Thermo Scientific, San Jose, CA). The peptide samples were loaded in 5_μL injections to an IonOpticks Aurora 0.075×250mm, 1.6 μm 120Å analytical column and column temperature was set to 50°C with a sonation oven. The flow rate was set at 400 nL/minute with solvent A (0.1% formic acid in water) and solvent B (0.1% formic acid and 80% acetonitrile) as the mobile phases. Separation was conducted using the following gradient: 3-19% B in 108 min; 19-29% B in 42 min; 29-41% B in 30 min. The full MS1 scan (m/z 350-1200) was performed on the Orbitrap Exploris with a resolution of 120,000. The FAIMS voltages were on with a FAIMS CV (V) set at −50. The RF Lens (%) was set to 40 and a custom automatic gain control (AGC) target was set with a normalized AGC target (%) set at 300. Monoisotopic precursor selection (MIPS) was enforced to filter for peptides with relaxed restrictions when too few precursors are found. Peptides bearing +2 - 6 charges were selected with an intensity threshold of 5e3. A custom dynamic exclusion mode was used with 60 s exclusion duration and isotopes were excluded. Data-dependent MS/MS was carried out with a 3 FAIMS CV loop (−50, −65, −80). MS/MS orbitrap resolving power was set to 60,000 with 2 m/z quadropole isolation. Top speed for data dependent acquisition within a cycle was set to 118 ms maximum injection time. The MS/MS mass tolerance was set to 10 ppm. Fragmentation of the selected peptides by higher energy collision dissociation (HCD) was done at 30% of normalized collision energy and a 2 (m/z) isolation window. The MS2 spectra were detected by defining first mass scan range as 120 m/z and the maximum injection time as 118 ms.

### Metabolite and Lipid Preparation and LC-MS/MS

The untargeted metabolomic approach used the high resolution Orbitrap Fusion Tribrid mass spectrometer (Thermo Fisher Scientific, Waltham, MA, USA) with Vanquish^™^ UHPLC liquid chromatography. An Accucore C18 (100 × 2.1) column was used for metabolites with solvent A (0.1% formic acid in water) and solution B (0.1% formic acid in acetonitrile). The column chamber temperature was to 55 °C. Pump flow rate was set to 0.45 mL/min. The LC gradient was set to 0 min: 1% of solvent B (i.e., 99% of solvent A), 5 min: 1% of B, 6 min: 40% of B, 7.5 min: 98% of B, 8.5 min: 98% of B, 9 min: 0.1% of B, 10 min stop run. To enhance identification, an Acquire X MSn data acquisition strategy was used which employs replicate injections for exhaustive sample interrogation and increases the number of compounds in the sample with distinguishable fragmentation spectra for identification. Electrospray ionization (ESI) was used in both positive and negative modes with a spray voltage for positive ions (V) = 3500 and a spray voltage for negative ions (V) = 2500. Sheath gas was set to 50, auxiliary gas was set at 1 and sweep gas was set to 1. The ion transfer tube temperature was set at 325 °C and the vaporizer temperature was set at 350 °C. Full MS1 used the Orbitrap mass analyzer (Thermo Fisher Scientific, Waltham, Massachusetts, USA) with a resolution of 120,000, scan range (m/z) of 55–550, MIT of 50, AGC target of 2e5, 1 microscan, and RF lens set to 50%. For untargeted lipidomics, a Vanquish HPLC-Q Exactive Plus system was used with an Acclaim C30 column (2.1 mm × 150 mm, 3μm). Solution A for lipids consisted of 0.1% formic acid, 10 mM ammonium formate, and 60% acetonitrile. Solution B for lipids consisted of 0.1% formic acid, 10 mM ammonium formate, and 90:10 acetonitrile: isopropyl alcohol. The column chamber temperature was set to 40 °C. Pump flow rate was set to 0.40 mL/min. The LC gradient was set to 0 min: 32% of solvent B (i.e., 68% of solvent A), 1.5 min: 45% of B, 5 min: 52% of B, 8 min: 58% of B, 11 min: 66% of B, 14 min: 70% of B, 18 min: 75% of B, 21 min: 97% of B, 26 min: 32% of B, 32 min stop run. The method for Q Exactive Plus mass spectrometer included a 32-min duration time, 10 s chromatogram peak width with full MS and ddMS2. Ion fragmentation was induced by HCD, with positive and negative polarity switching and a default charge state of 1. Full MS1 used the Orbitrap mass analyzer with a resolution of 70,000, 1 microscan, AGC target set to 1e6, and a scan range from 200 to 2000 m/z. The dd-MS2 scan used 1 microscan, resolution of 35,000, AGC target 5e5, MIT of 46 ms, loop count of 3, isolation window of 1.3 m/z, and a scan range of 200 to 2000 m/z for positive and negative polarity.

### Data analysis for Proteins, Metabolites, and Lipids

For LC-MS/MS proteomic data analysis, we used Proteome Discoverer^™^ 2.4 (Thermo Fisher Scientific, Waltham, MA, USA) with the search engine SEQUEST algorithm to process raw MS files. Spectra were searched using the TAIR10 protein database with the following parameters: 10 ppm mass tolerance for MS1 and 0.02 da as mass tolerance for MS2, two maximum missed tryptic cleavage sites, a fixed modification of carbamidomethylation (+57.021) on cysteine residues, dynamic modifications of oxidation of methionine (+15.996) and phosphorylation (+79.966) on tyrosine, serine, and threonine. Search results were filtered at 1% false discovery rate (FDR) and peptide confidence level was set for at least two unique peptides per protein for protein identification. Relative protein abundance in primed and control *dir1* and WS guard cell samples was measured using label-free quantification in Proteome Discoverer™ 2.4 (Thermo Scientific, Bremen, Germany). Proteins identified and quantified in all 4 out of 4 sample replicates were used. Peptides in mock and primed samples were quantified as area under the chromatogram peak. Peak areas were normalized by total protein amount. The average intensity of four primed *dir1* vs. four primed WS samples were compared as a ratio and two criteria were used to identify significantly altered proteins: (1) increase or decrease of 2-fold (primed *dir1*/primed WS), and (2) p-value from an unpaired Student’s t-test less than 0.05. For untargeted metabolomics, Compound Discover^™^ 3.0 Software (Thermo Scientific, Bremen, Germany) was used for data analyses. Raw files from four replicates of *dir1* primed and four replicates of WS primed guard cells were used as input. Spectra were processed by aligning retention times. Detected compounds were grouped and gaps filled using the gap filling node in Compound Discover that fills in missing peaks or peaks below the detection threshold for subsequent statistical analysis. Peak area was refined from normalize areas while marking background compounds. Compound identification included predicting compositions, searching mzCloud spectra database, and assigning compound annotations by searching ChemSpider, pathway mapping to KEGG pathways and to Metabolika pathways was included for functional analysis of the metabolites. The metabolites were scored by applying mzLogic and the best score was kept. Peak areas were normalized by the positive and negative mode internal standards (lidocaine and camphorsulfonic acid, respectively) added during sample preparation. For untargeted lipidomics data analyses, Lipid Search 4.1^™^ and Compound Discover^™^ 3.0 (Thermo Scientific, Bremen, Germany) were used. Raw files from three replicates of mock and three replicates of primed guard cells were uploaded Lipid Search 4.1^™^ for annotation of lipids found in all the samples. A mass list was generated for uploading to Compound Discover^™^ 3.0 Software. This mass list was used for compound identification along with predicted compositions, searching mzCloud spectra database, and assigning compound annotations by searching ChemSpider. Peak areas were normalized by median-based normalization. For both metabolomics and lipidomics, the average areas of four *dir1* primed vs. four WS primed metabolite samples were compared as a ratio and two criteria were used to determine significantly altered metabolites or lipids: (1) increase or decrease of 2-fold (*dir1* primed/WS primed), and (2) p-value from an unpaired Student’s t-test less than 0.05.

### Accession Numbers and Data Repository Information

The datasets presented in this study can be found in online repositories. The names of the repository/repositories and accession number(s) can be found below: All protein MS raw data and search results have been deposited to the ProteomeXchange Consortium via the PRIDE partner repository with the data set identifier PXD024991. All metabolite and lipid MS raw data and search results have been deposited to the MetaboLights data repository with the data set identifier MTBLS2614.

## Supplemental Data

The following supplemental materials are available.

**Supplemental Table S1.** Total proteins, metabolites, and lipids identified by LC-MS/MS from guard cells of WT and *dir1* knockout mutant under control and priming conditions.

**Supplemental Figure S1**. Singular enrichment analysis (SEA) for biological process using agriGO v2.0. shows pathway enrichment of proteins related to defense response, amino acid biosynthesis, and carbon metabolism in guard cells.

**Supplemental Figure S2**. Singular enrichment analysis (SEA) for cellular components using agriGO v2.0 shows pathway enrichment of plastid and cell-wall related proteins.

## ACKNOWLEDGMENTS

We thank Ms. Angelica Ortega and Mr. Ivan Grela their help with growth and maintenance of Arabidopsis plants. This material is based upon work supported by the National Science Foundation under Grant No. 1920420.

